# Efficient depletion of ribosomal RNA for RNA sequencing in planarians

**DOI:** 10.1101/670604

**Authors:** Iana V. Kim, Eric J. Ross, Sascha Dietrich, Kristina Döring, Alejandro Sánchez Alvarado, Claus-D. Kuhn

## Abstract

**Background:** The astounding regenerative abilities of planarian flatworms prompt a steadily growing interest in examining their molecular foundation. Planarian regeneration was found to require hundreds of genes and is hence a complex process. Thus, RNA interference followed by transcriptome-wide gene expression analysis by RNA-seq is a popular technique to study the impact of any particular planarian gene on regeneration. Typically, the removal of ribosomal RNA (rRNA) is the first step of all RNA-Seq library preparation protocols. To date, rRNA removal in planarians was primarily achieved by the enrichment of polyadenylated (poly(A)) transcripts. However, to better reflect transcriptome dynamics and to cover also non-poly(A) transcripts, a procedure for the targeted removal of rRNA in planarians is needed.

**Results:** In this study, we describe a workflow for the efficient depletion of rRNA in the planarian model species *S. mediterranea*. Our protocol is based on subtractive hybridization using organism-specific probes. Importantly, the designed probes also deplete rRNA of other freshwater triclad families, a fact that considerably broadens the applicability of our protocol. We tested our approach on total RNA isolated stem cells (termed neoblasts) of *S. mediterranea* and compared ribodepleted libraries with publicly available poly(A)-enriched ones. Overall, mRNA levels after ribodepletion were consisted with poly(A) libraries. However, ribodepleted libraries revealed higher transcript levels for transposable elements and histone mRNAs that remained underrepresented in poly(A) libraries. As neoblasts experience high transposon activity this suggests that ribodepleted libraries better reflect the transcriptional dynamics of planarian stem cells. Furthermore, the presented ribodepletion procedure was successfully expanded to the removal of ribosomal RNA from the gram-negative bacterium *Salmonella typhimurium*.

**Conclusions:** The ribodepletion protocol presented here ensures the efficient rRNA removal from low input total planarian RNA, which can be further processed for RNA-Seq applications. Resulting libraries contain less than 2% rRNA. Moreover, for a cost-effective and efficient removal of rRNA prior to sequencing applications our procedure might be adapted to any prokaryotic or eukaryotic species of choice.

## Background

Freshwater planarians of the species *Schmidtea mediterranea* are well known for their extraordinary ability to regenerate. This ability is supported by the presence of a large population of adult pluripotent stem cells, termed neoblasts [1]. Neoblasts are capable of producing all planarian cell types [2]. Moreover, they preserve their potency over the whole lifespan of the animal, which seems to be infinite [3]. Therefore, planarians embody an excellent model to study regeneration, aging and stem cell-based diseases. The phylum Platyhelminthes, to which *S. mediterranea* belongs to, includes multiple other members showing varying degrees of regenerative abilities. While some freshwater species (e.g. *Dugesia japonica* and *Polycelis nigra*) are capable to restore their body from any tiny piece [4, 5], others (*Procotyla fluviatilis*) have limited anterior regeneration abilities [6]. Altogether, the ability to regenerate is not solely based on the presence of pluripotent stem cells, but represents a complex interplay between different signaling pathways. The underlying changes in gene expression therefore need to be studied using transcriptome-wide techniques like RNA sequencing.

For any informative RNA-seq library preparation, ribosomal RNA, comprising >80% of total RNA, has to be removed. To achieve this goal two strategies can be pursued: either polyadenylated (poly(A)) RNA transcripts are enriched or rRNA is removed. Both approaches have advantages and limitations. On the one hand, the enrichment of poly(A) transcripts ensures better coverage of coding genes compared to ribodepleted samples, when sequenced to similar depth [7]. However, this advantage is outweighed by the loss of transcripts lacking poly(A) tails, which include preprocessed RNAs, a large share of all non-coding RNAs, such as enhancer RNAs and other long non-coding RNAs. In addition, long terminal repeat (LTR) retrotransposons and various intermediates of endonucleotic RNA degradation are lost during poly(A) selection [8–13]. Furthermore, most prokaryotic RNAs lack poly(A) tails, making rRNA depletion crucial for the study of bacterial transcriptome [14].

Here, we describe a probe-based subtractive hybridization workflow for rRNA depletion that efficiently removes planarian rRNA from total RNA. The protocol can be applied to input as low as 100 ng total RNA, which corresponds to 100,000 FACS-sorted planarian stem cells (X1 population) [15, 16]. Moreover, the DNA probes developed for *S. mediterranea* were successfully used for the removal of ribosomal RNA in related planarian species of the order Tricladida. The rRNA removal workflow presented here is also easily adapted to other organisms, as demonstrated by the removal of rRNA from *Salmonella typhimurium* total RNA using organism-specific probes.

## Results

### Development of an efficient rRNA depletion protocol for planarians

To deplete ribosomal RNA from planarian total RNA, we chose to develop a protocol based on the hybridization of rRNA-specific biotinylated DNA probes to ribosomal RNA and the capture of the resulting biotinylated rRNA-DNA hybrids by use of streptavidin-coated magnetic beads (Figure 1A). To that end, we synthesized a pool of 88 3’-biotinylated 40-nt long DNA oligonucleotide probes (siTOOLs Biotech, Martinsried, Germany). We chose probes with a length of 40 nucleotides since their melting temperature in RNA-DNA hybrids was shown to be 80±6.4 °C in the presence of 500 mM sodium ions [17]. This would allow probe annealing at 68°C in agreement with general probe hybridization temperatures [18]. The probes were devised in antisense orientation to the following planarian rRNA species: 28S, 18S type I and type II, 16S, 12S, 5S, 5.8S, internal transcribed spacer (ITS) 1 and ITS 2 (Supplementary Table1).

**Figure 1.**
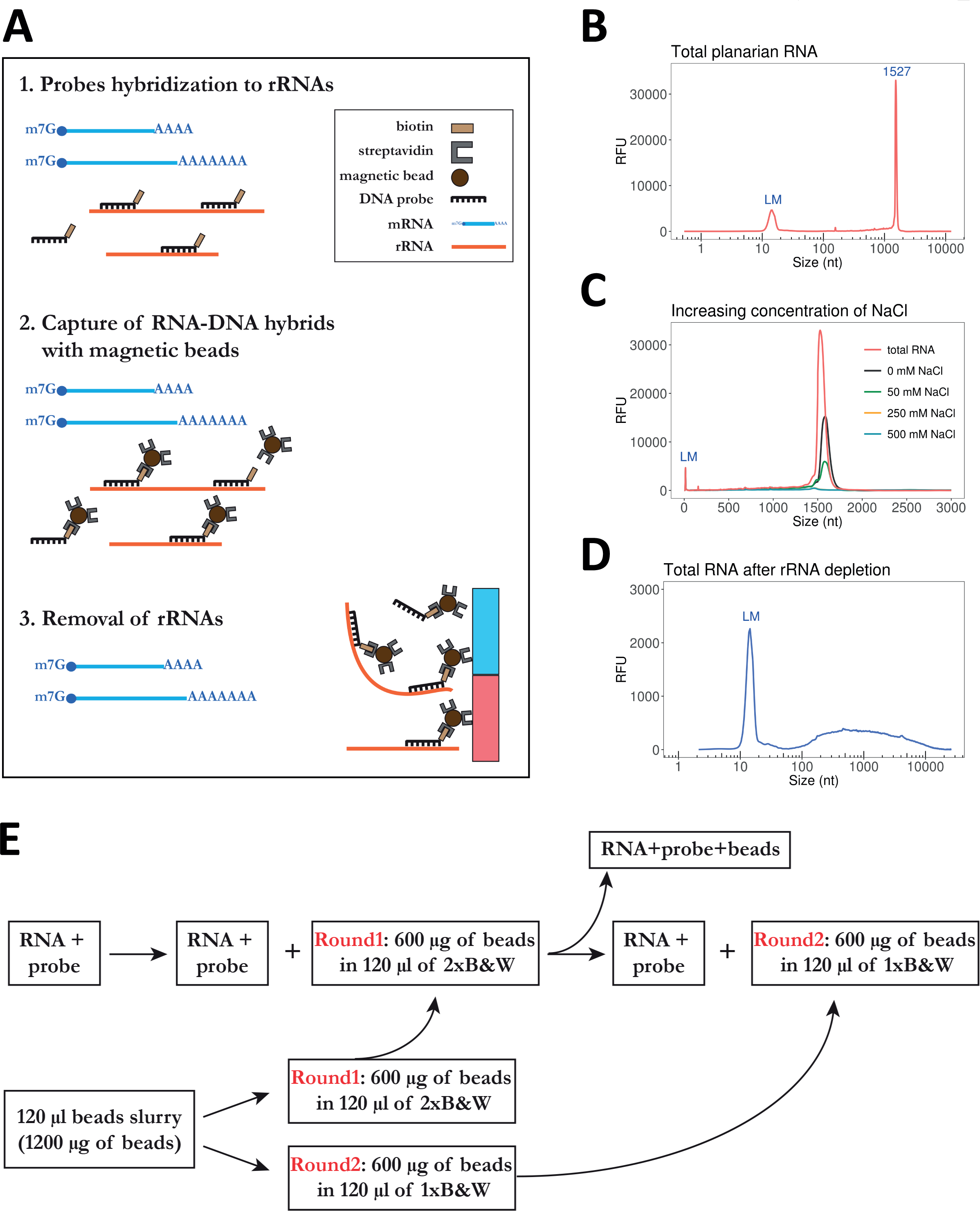
Efficiency of rRNA removal from total planarian RNA. **(A)** Schematic representation of rRNA depletion workflow. Biotinylated DNA probes are hybridized to rRNA, followed by subtraction of rRNA-DNA hybrids using streptavidin-labeled magnetic beads. **(B)** Separation profile of planarian total RNA. The large peak at 1527 nts corresponds to the co-migrating 18S rRNAs and the two fragments of processed 28S rRNA. LM stands for lower marker with the length of 15 nt (C) Increasing concentration of NaCl improves the efficiency of rRNA removal. (D) Total planarian RNA after rRNA depletion. (E) Removal of rRNA-DNA hybrids was performed in two consecutive steps using streptavidin-coated magnetic beads resuspended in 2× of 1× B&W buffer.

To assess RNA quality and the efficiency of rRNA removal we used capillary electrophoresis (Fragment Analyzer, Agilent). The separation profile of total planarian RNA only shows a single rRNA peak at about 1500 nucleotides (nts) (Figure 1B). This single rRNA peak is the result of the 28S rRNA being processed into two fragments that co-migrate with the peak for 18S rRNA [19]. Planarian 28S rRNA processing usually entails the removal of a short sequence located in the D7a expansion segment of 28S rRNA. The length of the removed fragment thereby varies from 4 nts to 350 nts between species (e.g. for *Dugesia japonica* 42 nts are removed) [19]. Intriguingly, a similar rRNA maturation process was observed in particular protostomes, in insects such as *D. melanogaster* and in other platyhelminthes [19–21]. In addition to this 28S rRNA maturation phenomenon, *S. Mediterranea* possesses two 18S rDNA copies that differ in about 8% or their sequence. However, only 18S rRNA type I was reported to be functional and predominantly transcribed [22, 23].

As a first step during rRNA removal all 88 DNA probes were annealed to total planarian RNA. Since RNA molecules are negatively charged, the presence of cations facilitates the annealing of probes to RNA by reducing the repulsion of phosphate groups [24, 25]. Although Mg^2+^ions are most effective in stabilizing the tertiary structure of RNA and in promoting the formation of DNA-RNA hybrids, they are also cofactors for multiple RNases [26] and hence should not be included during ribodepletion. Therefore, we tested several hybridization buffers with varying concentrations of sodium ions (Figure 1C). In the absence of sodium ions we only accomplished the incomplete removal of rRNA. However, sodium concentrations >250 mM in the hybridization buffer led to the complete depletion of rRNA from planarian total RNA (Figure 1C, 1D). Thus, optimal rRNA removal requires the presence of >250 mM NaCl in the hybridization buffer. As we obtained the most consistent results in the presence of 500 mM NaCl, we decided to utilize this salt concentration in our procedure (Figure 1D).

### Detailed rRNA depletion workflow

#### Required buffers

Hybridization buffer (20 mM Tris-HCl (pH 8.0), 1 M NaCl, 2 mM EDTA)

Solution A (100 mM NaOH, 50 mM NaCl, DEPC-treated)

Solution B (100 mM NaCl, DEPC-treated)

2×B&W (Binding&Washing) buffer (10 mM Tris-HCl (pH 7.5), 1 mM EDTA, 2 M NaCl)

Dilution buffer (10 mM Tris-HCl (pH 7.5), 200 mM NaCl, 1 mM EDTA)

#### Protocol

1. RNA input The following protocol efficiently depletes ribosomal RNA from 100 up to 1.5 µg of total RNA (Figure 1E). The procedure can be scaled up for higher RNA input.
2. Hybridization of biotinylated DNA oligonucleotides (40-mers) to ribosomal RNA.
  a. For oligonucleotide annealing the following reaction is set up: 10 µl hybridization buffer 10 µl RNA input (1 µg) 1 µl 100 µM biotinylated DNA 40-mers probes
  b. Gently mix the solution by pipetting and incubate at 68 °C for 10 min.
  c. Immediately transfer the tubes to 37 °C for 30 min.
3. Prepare Dynabeads MyOne Streptavidin C1 (Invitrogen) according to the manufacturer’s instruction as follows:
  a. For each sample use 120 µl (10 µg/µl) of beads slurry.
  b. Wash the beads twice with an equal volume (or at least 1 ml) of Solution A. Add Solution A and incubate the mixture for 2 min. Then, place the tube on a magnet for 1 min and discard the supernatant.
  c. Wash the beads once in Solution B. Split the washed beads into two separate tubes for two rounds of subtractive rRNA depletion (Round1 and Round2). Place the beads on a magnet for 1 min and discard Solution B.
  d. Resuspend the Round1 beads in 2xB&W buffer to a final concentration of 5 µg/µl (twice the original volume). The Round1 beads will be used during the first round of rRNA depletion. For the second round of depletion, resuspend the Round2 beads to a final concentration of 5 µg/µl in 1×&W buffer. The Round2 beads will be used in a second depletion step. Keep it at 37 °C until required.
4. Capture of DNA-RNA hybrids using magnetic beads (step 2).
  a. Briefly spin down the tubes containing total RNA and probes. Add the following: 100 µl dilution buffer 120 µl washed magnetic beads (5 µg/µl) in 2xB&W (Round1). Resuspend by pipetting up and down ten times. The final concentration of NaCl during this step is 1 M. Incubate the solution at 37 °C for 15 min. Gently mix the sample occasionally by tapping.
  b. Place on magnet for 2 min. Carefully remove the supernatant and add the additional 120 µl of washed magnetic beads in 1xB&W (Round2). Incubate the mixture at 37°C for 15 min with occasional gentle tapping.
  c. Place on magnet for 2 min. Carefully transfer the supernatant into a new tube and place it on magnet for another 1 min to remove all traces of magnetic beads from the sample.
  d. Transfer the supernatant into a fresh tube.
5. Use the RNA Clean µ Concentrator-5 kit (Zymo Research) to concentrate the ribodepleted samples, to carry out size selection and to digest any remaining DNA using DNase I treatment as described [27].

### Ribosomal RNA depletion in planarian species related to *S. mediterranea*

Ribosomal DNA genes are among the most conserved sequences in the kingdom of life. They are present in all organisms and are widely used for the construction of phylogenetic trees [28]. The latter is possible because of the low rate of nucleotide substitutions in rRNA sequences (about 1 - 2% substitutions occur per 50 million years based on bacterial 16S rRNA) [29]. The divergence of 18S rRNA sequence between different families of freshwater planarians lays in the range of 6 – 8%, while interspecies diversity does not exceed 4% [23]. Therefore, low rRNA divergence between taxa can be exploited for the design of universal probes for rRNA depletion in different organisms. To assess the specificity and universal applicability of our DNA probes, we depleted rRNA in flatworm species of the order Tricladida, all related to S. mediterranea (Figure 2A). Total RNA separation profiles were analyzed before and after rRNA depletion of six planarian species from three different families. Two of these, *Dugesia japonica* and *Cura pinguis* belong to the same family as *S. mediterranea*, the Dugesiidae family. In addition, we examined three species from the family Planariidae (*Planaria torva, Polycelis nigra* and *Polycelis tenuis*) and one species from the genus *Camerata* of Uteriporidae (subfamily Uteriporinae). For all tested species our DNA probes proved efficient for the complete removal of rRNA, which migrated close to 2000 nts on all electropherograms (Figure 2B). Thus, the probes presented here can be utilized for the removal of ribosomal RNA in a multitude of planarian species and may even be generally applicable to all studied planarian species.

**Figure 2.**
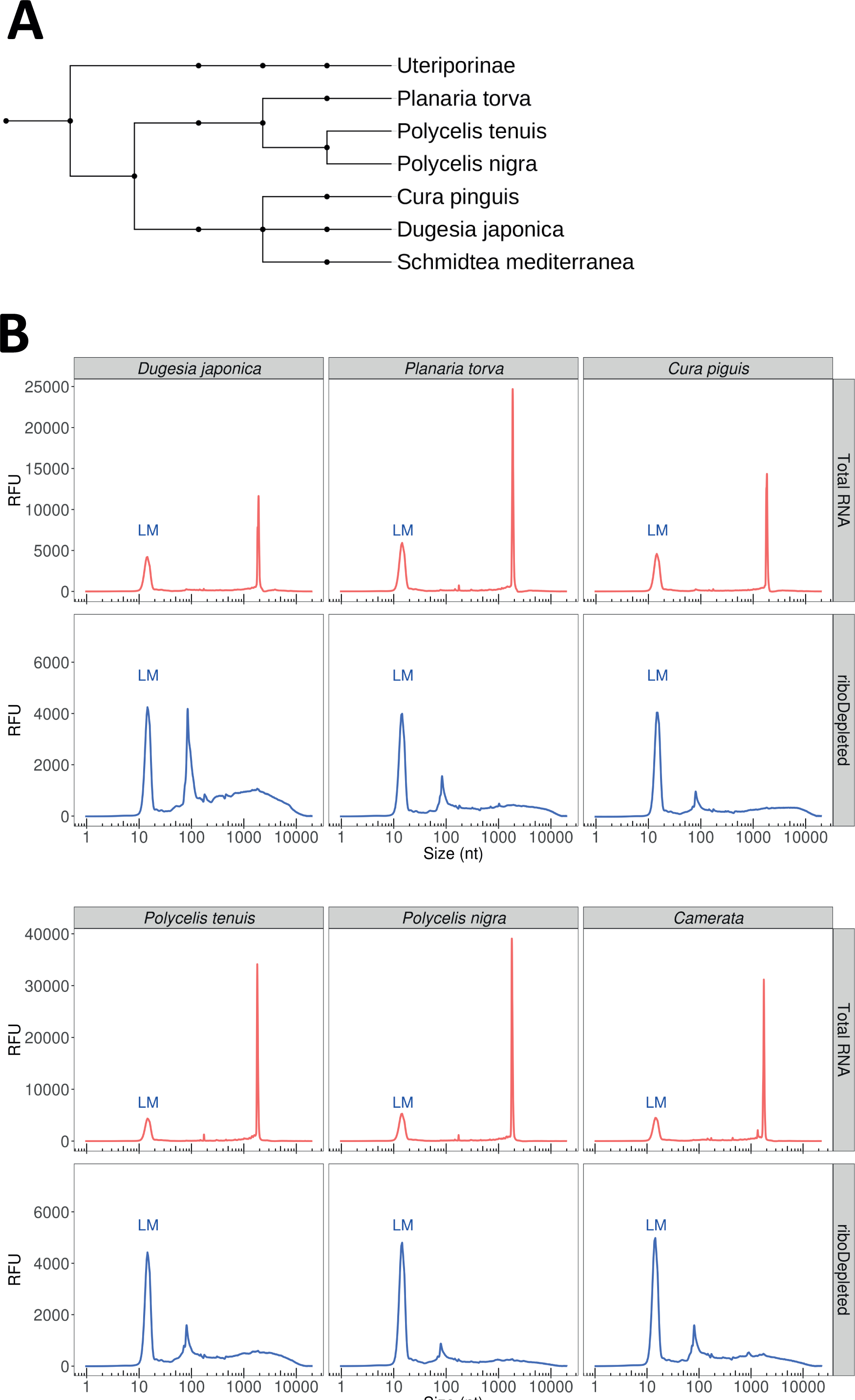
Probes developed for *S. mediterranea* efficiently remove rRNA of other freshwater triclads. **(A)** Phylogenetic tree showing the taxonomic position of the analyzed planarian species. **(B)** Separation profile of total RNA before and after rRNA depletion. In all species analyzed the 28S rRNA undergoes the “gap deletion” maturation that result in two co-migrating fragments. Both 28S fragments co-migrate with 18S rRNA, resulting in a single rRNA peak. The peak at about 100 nts represents a variety of small RNAs (5S and 5.8S rRNA, tRNAs, and other small RNA fragments) that evaded size selection by the RNA Clean µ Concentrator-5 kit (Zymo Research).

### Comparison of RNA-Seq libraries prepared by ribodepletion or poly(A) selection

To assess the efficiency of rRNA removal and the specificity of our DNA probes, we prepared and analyzed RNA-Seq libraries from ribodepleted planarian total RNA from *S. mediterranea*. Total RNA was extracted from 100,000 FACS-sorted planarian neoblasts, resulting in 70 – 100 ng of input RNA. RNA-Seq libraries were prepared and sequenced as described [27] following 15 cycles of PCR amplification. The subsequent analysis of sequenced libraries confirmed the efficient removal of rRNAs. Less than 2% of total sequenced reads constituted ribosomal RNA (Figure 3A). Next, we compared our rRNA-depleted libraries with three publicly available planarian poly(A) enriched RNA-Seq datasets (poly(A) libraries) [30–32]. In case publicly available libraries were sequenced in paired-end mode, we analyzed only the first read of every pair to minimize the technical variation between libraries [33]. As shown in Figure 3A, the ribodepleted libraries contained significantly less rRNA compared to all poly(A) enriched ones. Interestingly, the major rRNA species that remained after poly(A) selection was mitochondrial 16S rRNA (Figure 3B). Although the planarian genome has a high A-T content (> 70%) [34], we could not attribute the overrepresentation of 16S rRNA in poly(A) libraries to a high frequency or longer stretches of A nucleotides as compared to other rRNA species (Figure 3C). Moreover, using publicly available planarian poly(A)-position profiling by sequencing (3P-Seq) libraries [35] which allow the identification of 3’-ends of polyadenylated RNAs, no polyadenylation sites were detected on 16S rRNA. Therefore, we speculate that upon folding of the 16S rRNA stretches of A nucleotides become exposed and facilitate the interaction with oligo-dT beads during transcript poly(A) selection.

**Figure 3.**
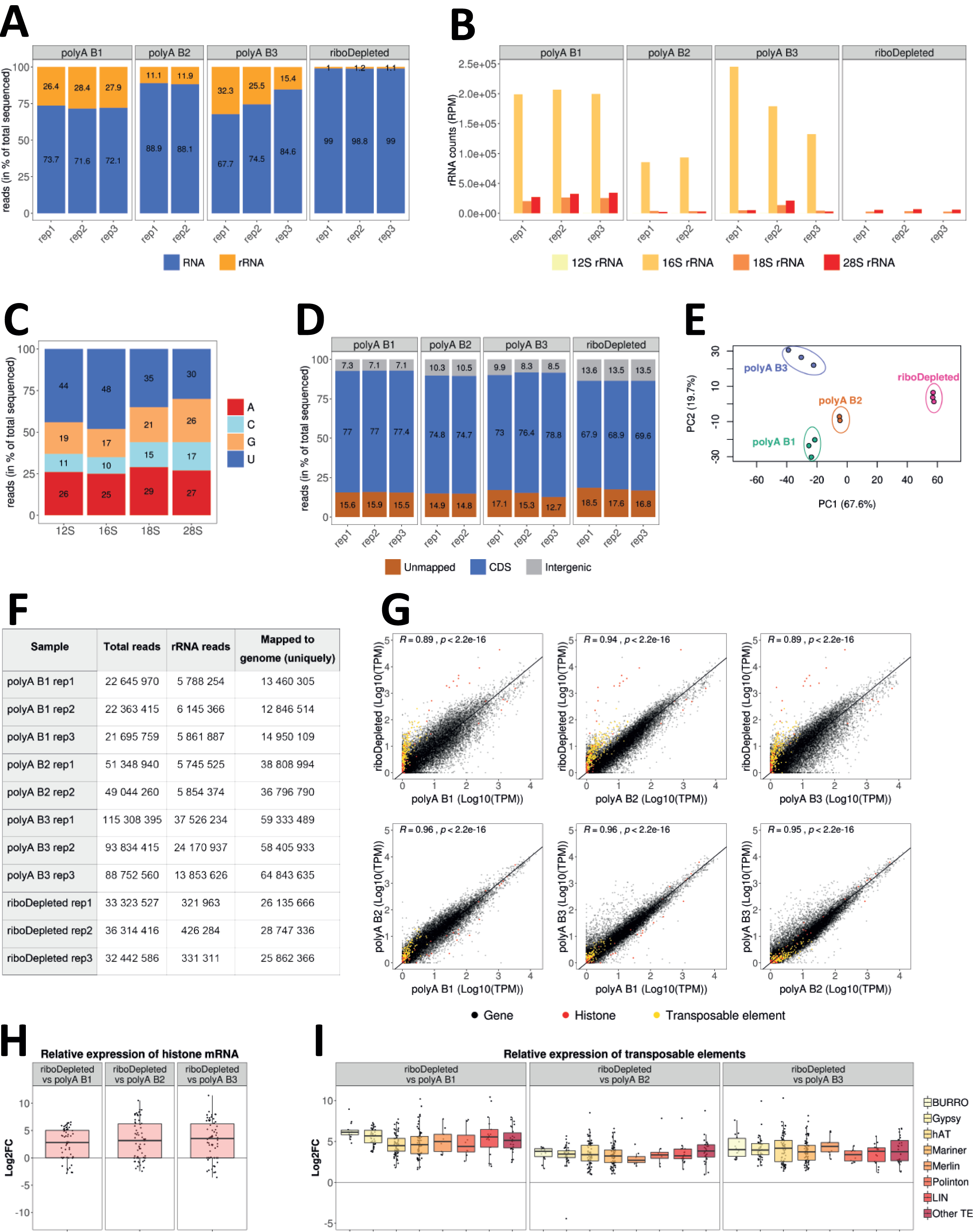
Comparison of rRNA-depleted and poly(A)-enriched planarian RNA-Seq libraries. **(A)** Percentage of rRNAs reads in the sequenced libraries prepared from rRNA-depleted or poly(A)-enriched RNA. **(B)** Species of rRNAs remaining in the final sequenced libraries. **(C)** Nucleotide content of planarian rRNA. **(D)** Percentage of sequenced reads mapped to coding (CDS) and intergenic regions in the planarian genome. **(E)** Principal component analysis (PCA) biplot of log2 expression data for coding genes reveals distinct clustering of all analyzed RNA-Seq experiments. **(F)** Sequencing depth and number of reads mapped to the planarian genome in analyzed rRNA-depleted and poly(A)-enriched samples. **(G)** Comparison of gene expression in transcripts per million (TPM) between planarian ribodepleted and poly(A)-enriched (polyA) RNA-Seq data. The Pearson’s correlation coefficient is indicated. **(H)** Increased representation of histone mRNAs in ribodepleted libraries. **(I)** Boxplot of log2 fold changes of the expression of transposable elements between ribodepleted and poly(A)-enriched libraries.

We next assigned the analyzed datasets to the planarian genome. In ribodepleted libraries more than 13% of all mapped reads were assigned to intergenic regions, compared to 7% – 10.5% for poly(A)-enriched ones (Figure 3D). In addition, the percentage of unmapped reads was higher in ribodepleted libraries and constituted about 17.6%, which is on average 2.4% more than in poly(A) datasets. We speculate that for ribodepleted libraries the proportion of reads mapping to intergenic regions will increase in future once complete assemblies of the planarian genome are available. Currently, the planarian genome assembly consists of 481 scaffolds [34]. To detect gene expression variabilities between the analyzed libraries, we performed principal component analysis for the clustering of gene expression data. Although all poly(A) selected libraries were grouped closer together along the PC1 scale, all four analyzed datasets appeared as separated clusters. This indicates the presence of considerable variation even amongst different batches of poly(A) libraries (Figures 3E). One possible source of such variation might be the sequencing depth of the analyzed libraries, which varied considerably from 13 to 64 million of mapped reads (Figure 3F).

Next, to estimate the correlation between ribodepleted and poly(A) libraries, we calculated their Pearson correlation coefficients (Figure 3G). We found the highest Pearson correlation between ribodepleted libraries and polyA B2 samples (R=0.94, p < 2.2e-16) (Figure 3F). This could be due to their similar sequencing depth compared to the other polyA libraries. The transcripts whose abundance was most significantly affected by poly(A) selection were found to be histone mRNAs that are known to lack polyA tails (Figure 3G, 3H) [36]. Their expression level appeared to be 8 – 10 fold higher in our ribodepleted libraries. Moreover, in the ribodepleted libraries we detected significantly higher expression levels for transposable elements (Figure 3G, 3I). Out of 316 planarian transposable element families [37] 254 were on average upregulated 5.2, 3.5 and 4.0 fold as compared to polyA B1, polyA B2 and polyA B3 libraries, respectively (Figure 3I). Our ribodepleted libraries revealed that Burro elements, giant retroelements found in planarian genome [34], gypsy retrotransposons, hAT and Mariner/Tc1 DNA transposons are the most active transposable elements in planarian stem cells. Although some of transposable elements are polyadenylated, long-terminal repeat elements (LTRs) lack poly(A)-tails [38]. This renders their detection in poly(A)-enriched sample non-quantitative. Last, our rRNA depletion workflow also allows for the detection of transposable element degradation products that were generated by PIWI proteins guided by specific piRNAs, an abundant class of small RNAs in planarians [39, 40].

### Non-specific depletion of coding transcripts in ribodepleted libraries

In using custom ribodepletion probes, our major concern was that the utilized probes would lead to unspecific co-depletion of planarian coding transcripts. To exclude this possibility, we first mapped our pool of 88 DNA probes in antisense orientation to the planarian transcriptome allowing up to 8 mismatches and gaps of up to 3 nts. This mapping strategy requires at least 75% of a DNA probe to anneal to its RNA target. It resulted in only 11 planarian genes to be potentially recognized by 20 DNA probes from our oligonucleotide pool. Next, we carried out a differential expression analysis of these 11 potentially targeted transcripts between the ribodepleted libraries and poly(A)-selected ones. The analysis revealed that 9 out of 11 potential targets were downregulated at least 1-fold in at least two poly(A) experiments (Figure 4A). As the abundance of three transcripts (SMESG000014330.1 (rhodopsin-like orphan gpcr [41]), SMESG000068163.1 and SMESG000069530.1 (both without annotation)) was very low in all polyA libraries (<0.6 transcripts per million (TPM)) we did not consider these any further. However, we found the remaining six transcripts to be significantly downregulated in ribodepleted libraries. For three of these targeted genes (SMESG000067473.1, SMESG000021061.1 and SMESG000044545.1) the probes map in regions that display significant RNA-seq coverage (Figure 4B, Supplemental Figs. S1A, S1B). Therefore, their lower expression values in ribodepleted libraries is likely attributed to probe targeting. Intriguingly, for the remaining three targets (SMESG000066644.1, SMESG000043656.1 and SMESG000022863.1 annotated as RPL26 (ribosomal protein L26), COX11 (cytochrome c oxidase copper chaperone) and an unknown transcript, respectively) the probes were predicted to map to loci that do not exhibit RNA-seq coverage (Figure 4C, Supplemental Figs. S1C, S1D). The likely reason for this is inaccurate gene annotation. Alternatively, target regions might represent repetitive, multimapping sequences, which we excluded during read mapping. Taken together, our off-target analysis revealed that a maximum of 11 genes might be affected by our rRNA removal procedure - a very low number that underscores the specificity and efficiency of our depletion protocol.

**Figure 4.**
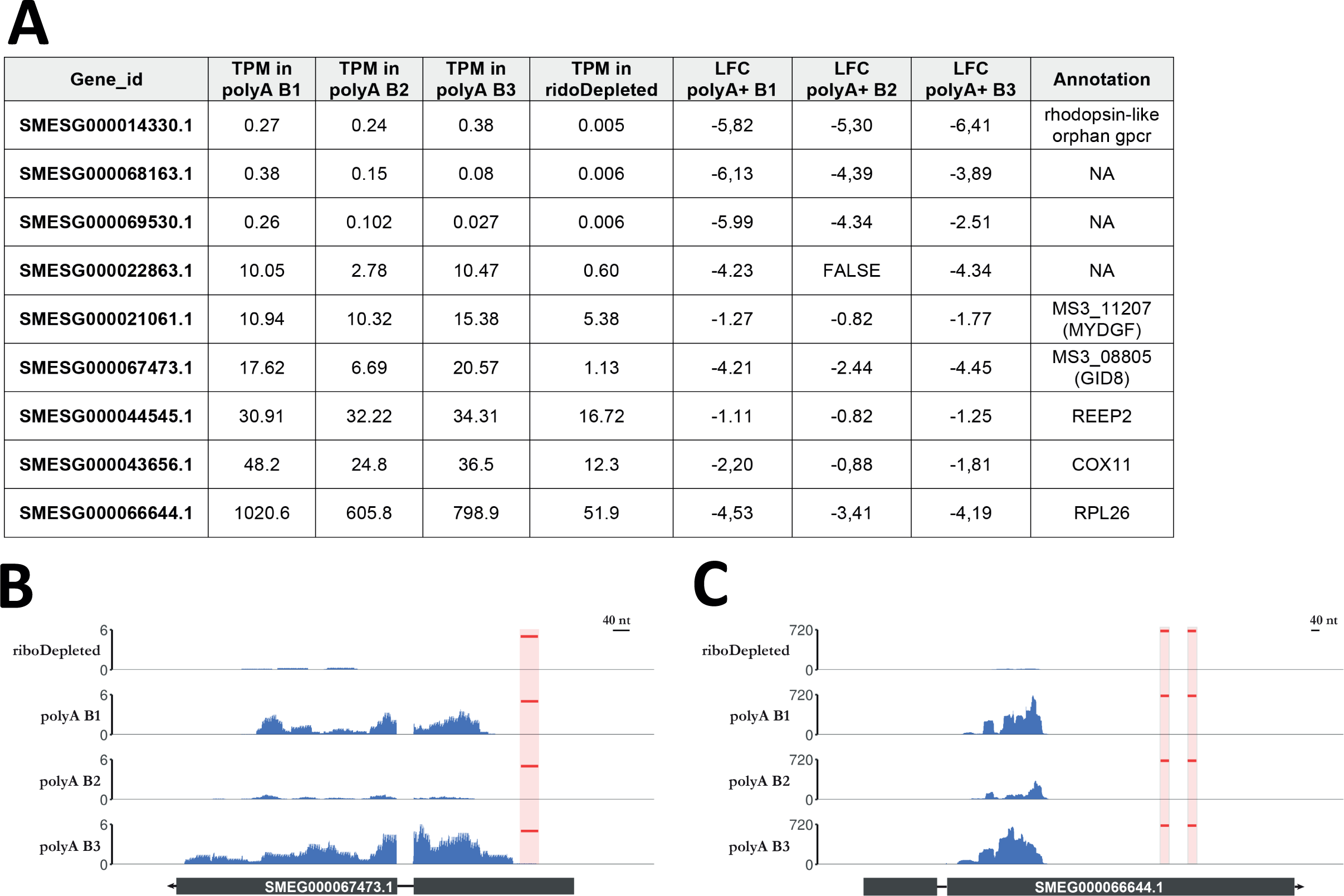
Off-target analysis of DNA probes used for rRNA depletion. **(A)** Expression level of nine transcripts targeted by designed probes. **(B)** RNA-Seq coverage profile for SMESG000067473.1 in rRNA depleted (ribodepleted) and poly(A) enriched (polyA B1, polyA B2, polyA B3) libraries. The position of antisense probes mapping to the transcripts is marked in red. **(C)** The same as (B) for SMESG000066644.1.

### Applicability of the described ribodepletion method to other organisms

To demonstrate the applicability of the developed rRNA workflow to other organisms, we employed our protocol to the depletion of ribosomal RNA from *Salmonella typhimurium* using a pool of organism-specific DNA probes (riboPOOL) developed by siTOOLs Biotech (Martinsried, Germany) (Figure 5A). We compared the libraries resulting from the application of our newly developed procedure to the established rRNA depletion workflow that utilizes the Ribo-Zero rRNA Removal Kit (Bacteria) from Illumina. Removal of rRNA from a *S. typhimurium* sample using riboPOOL probes was comparably successful as a depletion reaction using Ribo-Zero, leaving as low as 3.4% rRNA in the final library (Figure 5A). Moreover, an overall comparison of gene expression levels showed a high correlation (Pearson correlation > 0.98) between riboPOOL depleted libraries and libraries prepared with the Ribo-Zero kit (Figure 5B). Taken together, the rRNA depletion workflow described in this manuscript is robust and easily applicable to any bacterial and eukaryotic species of choice utilizing organism-specific probes.

**Figure 5.**
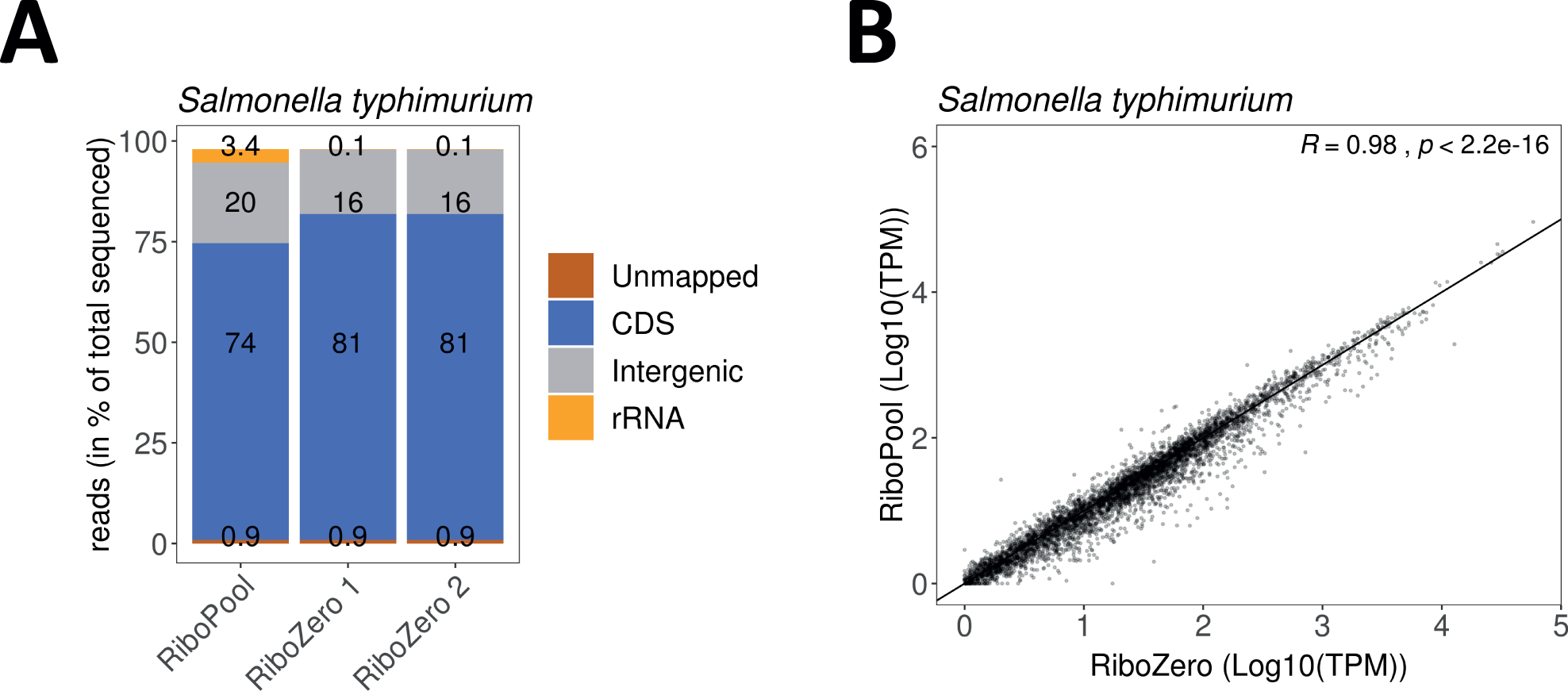
Application of the developed rRNA workflow to other species using organism-specific probes. **(A)** Percentage of rRNAs in *Salmonella typhimurium* sequenced libraries prepared using our developed rRNA depletion workflow with organism-specific RiboPool probes or the commercial Ribo-Zero kit. **(B)** Scatter plot showing the transcript abundance (transcript per million (TPM)) between rRNA depleted libraries using our developed workflow or the Ribo-Zero commercial kit. The Pearson’s correlation coefficient is indicated.

## Discussion

For samples from typical model organisms, such as human, mouse and rat, there are numerous commercial kits available for the removal of rRNA, e.g. NEBNext from New England Biolabs, RiboGone from Takara and RiboCop from Lexogen. This also applies to typical gram-positive and gram-negative bacteria (MICROBExpress from Thermofisher and Ribominus from Invitrogen). Moreover, these kits can be utilized with a certain degree of compatibility for the depletion of rRNA in organisms of distant phylogenetic groups (e.g. RiboMinus Eukaryote Kit for RNA-Seq, Invitrogen). However, as the breadth of molecularly tractable organisms increases in the past decade, the necessity to develop organism-specific rRNA depletion techniques rises as well [42–44]. To date, such custom protocols were either based on rRNA removal by biotinylated antisense probes along with streptavidin-coated magnetic beads or they were based on the digestion of DNA-RNA hybrids with RNase H [14, 45–47].

In this study, we describe a novel rRNA depletion workflow for the planarian flatworm *S. mediterranea*. Our protocol is based on the hybridization of biotinylated DNA probes to planarian rRNA followed by the subsequent removal of the resulting rRNA-DNA hybrids with streptavidin-labeled magnetic beads. A comparative analysis between ribodepleted and poly(A)-selected libraries in planarians revealed that our protocol retains all information present in poly(A) selected libraries. Over and above, we found ribodepleted libraries to contain additional information on histone mRNAs, transposable elements (of which many do not have polyA tails) and transcriptional intermediates. This highlights that ribodepletion better retains the dynamic information of transcriptomes. As more and more co-translational decay pathways are being discovered [48], understanding the dynamic of RNA degradation will become more important in the near future. Moreover, by successfully depleting rRNA from other freshwater triclad species, we could demonstrate the versatility of the DNA probes designed for *S. mediterranea*. Last, we validated the efficiency of the developed workflow by removal of rRNA in the gram-negative bacterium *S. typhimurium*. Therefore, the proposed workflow serves as an efficient and cost-effective method for rRNA depletion in any organism of interest.

## Conclusions

This study describes an rRNA depletion workflow for the planarian model system *S. mediterranea* and related freshwater triclads. It is based on the hybridization of 40-mer biotinylated DNA oligos to ribosomal RNA followed by the subtraction of formed DNA-RNA hybrids. The protocol is very robust and ensures the efficient removal of rRNA even from low input total RNA. Moreover, we suggest the general applicability of the presented workflow to any prokaryotic or eukaryotic organisms by using organism-specific pools of probes.

## Materials and Methods

### Ribosomal RNA depletion

Ribosomal RNA depletion was conducted as described in the result section. To evaluate Fragment analyzer separation profiles, planarian total RNA (1000 ng each sample) was subjected to rRNA depletion using varying concentrations of NaCl (0 mM, 50 mM, 250 mM, 500 mM) in the hybridization buffer.

### Phylogenetic tree

The phylogenetic tree was constructed using NCBI taxonomic names at phyloT (https://phylot.biobyte.de). The tree was visualized using the Interactive Tree of Life (iToL) tool [49].

### Planarian rRNA-depleted RNA-Seq dataset

Raw sequencing reads for planarian rRNA-depleted dataset were downloaded from the project GSE122199 (GSM3460490, GSM3460491, GSM3460492). The libraries were prepared as described [40]. Briefly, planarian rRNA depleted RNA-Seq libraries were prepared from 100,000 FACS-sorted planarian X1 cells as described [27] and sequenced on an Illumina Next-Seq 500 platform (single-end, 75 bp).

### Publicly available RNA-Seq datasets

Raw sequencing reads for all datasets were downloaded from the Sequence read archive (SRA). Planarian polyA B1 rep1, polyA B1 rep2, polyA B1 rep3 correspond to SRR2407875, SRR2407876, and SRR2407877, respectively, from the Bioproject PRJNA296017 (GEO: GSE73027) [30].

Planarian polyA B2 rep1, polyA B2 rep2 samples correspond to SRR4068859, SRR4068860 from the Bioproject PRJNA338115 [32]. Planarian polyA B3 rep1, polyA B3 rep2, polyA B3 rep3 correspond to SRR7070906, SRR7070907, SRR7070908, respectively, (PRJNA397855) [31]. Only first read of the pair was analyzed for polyA B2 and polyA B3 from Bioprojects PRJNA338115 and PRJNA397855.

### *Salmonella typhimurium* SL1344 RNA Seq datasets

In total, four samples were sequenced for *Salmonella typhimurium* SL1344 by the IMGM Laboratories GmbH (Martinsried, Germany) on an Illumina NextSeq 500 platform (single-end, 75bp). Of these sequenced samples, one sample was of untreated total RNA, two samples were of RiboZero and one of RiboPools treated total RNA. The sequencing data are available at NCBI Gene Expression Omnibus (http://www.ncbi.nlm.nih.gov/geo) under the accession number GSE132630.

### Processing of RNA-Seq libraries

Planarian RNA-Seq data were processed as follows: Reads after removal of 3’-adapters and quality filtering with Trimmomatic (0.36) [50] were trimmed to a length of 50 nt. For libraries sequenced in pair-end mode, only the first read of a pair was considered for the analysis. Next, sequences mapped to planarian rRNAs were removed with SortMeRNA [51]. Reads were assigned to the reference genome version SMESG.1 [34] or consensus transposable element sequences [37] in strand-specific mode and quantified with Kallisto [52] with “--single -l 350 -s 30 -b 30”. Differential gene expression analysis was performed with DeSeq2 [53]. To annotate RNA-Seq reads to coding regions (CDS), reads were mapped to the planarian genome using STAR [54] with the following setting: --quantMode TranscriptomeSam --outFilterMultimapNmax 1.

RNA sequencing data from *Salmonella typhimurium* SL1344 were processed with READemption 0.4.3 using default parameters [55]. Sequenced reads were mapped to the RefSeq genome version NC_016810.1 and plasmids NC_017718.1, NC_017719.1, NC_017720.1.

### Analysis of DNA probe specificity

DNA probe sequences were mapped to the planarian transcriptome SMEST.1 [56] using the BURST aligner (v0.99.7LL; DB15) [57] with the following settings “-fr -i .80 -m FORAGE”. Only sequences that mapped to genes in antisense orientation with no more than 8 mismatches were considered as potential probe targets.

## Acknowledgements

We thank Jochen Rink, Mario Ivankovic and Miquel Vila Farré from the Max Planck Institute of Molecular Cell Biology and Genetics in Dresden for generously providing various flatworm species. We thank Jens Hör from the Institute for Molecular Infection Biology, University of Würzburg, for providing the *Salmonella* RNA and IMGM Laboratories for library preparation and sequencing of *Salmonella* samples. We also acknowledge the KeyLab Genomics & Bioinformatics at the University of Bayreuth for Fragment Analyzer (Agilent) measurements. This work was supported by the Elite Network of Bavaria, the University of Bayreuth, the Paul Ehrlich and Ludwig Darmstaedter Prize for Young Researchers (to C.D.K) and the IZKF at the University of Würzburg (project Z-6). A.S.A. is an investigator of the Howard Hughes Medical Institute and the Stowers Institute for Medical Research.

## Authors’ contributions

I.V.K. and C.D.K. conceived and designed the study; I.V.K. and K.D. acquired data; E.J.R. assembled planarian rRNA sequences; I.V.K., E.J.R. and S.D. performed computation analyses; A.S.A. and C.D.K. supervised the study; I.V.K. and C.D.K. drafted the manuscript with input from all authors.

---

**Supplemental Figure S1.**
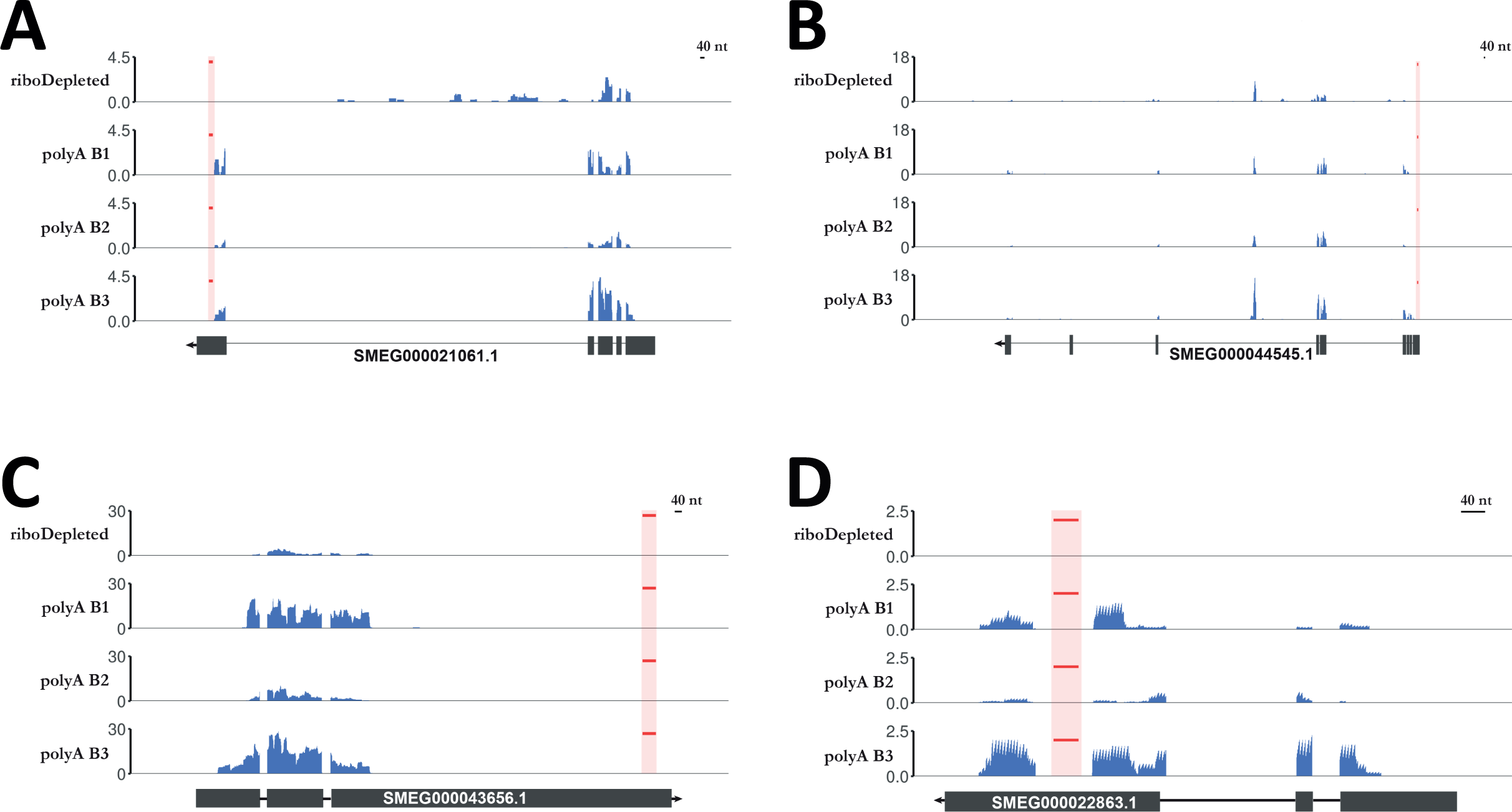
**(A)-(D)** RNA-Seq coverage profile for genes potentially targeted by designed probes in rRNA depleted (ribodepleted) and poly(A) enriched (polyA B1, polyA B2, polyA B3) libraries. The position of antisense probes mapping to the transcripts is marked in red.

